# The combined toxicity test of polyester and tetra ethylene glycol on *Daphnia magna*

**DOI:** 10.1101/2021.07.24.453624

**Authors:** Yeonjo Jung

## Abstract

The combined toxicity test of polyester and tetra ethylene glycol on *Daphnia magna*. Globally, wide use of plastics and its increased production has led to a mounting amount of plastic waste entering the natural ecosystems. Due to their small size, plastic particles might be ingested by organisms at the lower end of the food chain and can be transferred by feeding to top consumers. Consequently, plastic pollution in aquatic environments and its potential impact on aquatic life has recently been recognized as an issue of considerable concern for ecosystem. I produced microplastics from 100% polyester thread from Houjix, cut it finely and used a dissecting needle to cut into a size of 5mm or less through a microscope. I also included ethylene glycol to investigate its toxic effects on *D. magna*. Since my aim was also to compare the toxicity effect of both chemicals, I used different concentrations individually and then in combinations to determine the potential toxic effects of polyester and tetra ethylene glycol (combined and separately) on the life (survival, death) of *D. magna*.

Microplastics from 100% polyester thread from Houjix were produced, into a size of 5mm or less through a microscope. Tetra ethylene glycol was also used to investigate its toxic effects on *D. magna.* The toxicity effect of both chemicals using different concentrations individually and then in combinations were employed to determine the potential toxic effects of polyester and tetra ethylene glycol (combined and separately) on the life of *D. magna*. The study exhibited that the IC50 of TEG was higher as compared to polyester which suggests that polyester was more adverse than TEG. Moreover, when TEG and polyester were treated in combination, IC50 value was lower (Figure 3) than the IC50 value of TEG and polyester separately. In other words, the TEG and polyester in combination exhibited the lowest IC50 value in this study. These results suggest that TEG and polyester in combination had adverse effects on the growth and development of *D. magna*

## 1. Introduction

Plastics are of significant benefit for modern-day society because of their easy manufacturing, low price, and practical function in a variety of products. Wide use of plastics at global scale and its increased production has led to a mounting amount of plastic waste entering the natural ecosystems. It is estimated that between 1.15 and 2.41 million tons of plastic waste enter the oceans every year (Lebreton 2017). Plastics can degrade into a wide range of sizes, including micro- (<5 mm) and nano-sized (<100 nm) particles. Furthermore, pollution by micro- and nanoplastics constitutes a potential threat to aquatic ecosystems. Due to their small size, plastic particles might be ingested by organisms at the lower end of the food chain and can be transferred by feeding to top consumers. Several studies have shown that plastic particles of various sizes can be ingested by aquatic organisms causing tissue damage or even death. Therefore, plastic pollution (polyvinyl chloride, polyethylene, polyethylene terephthalate, polypropylene, polyvinyl alcohol) in aquatic environments and its potential impact on aquatic life has recently been recognized as an issue of considerable concern for society, as well as for ecosystem functioning (Chae 2018, Jeon 2018).

At the same time, organic solvents used in dry cleaning also cause toxicity and environmental pollution problems. Irregular use of tetra-ethylene glycol (an organic solvent) used for production of polyethylene terephthalate (PET), in dry cleaning, deicing fluids and for the antifreeze mixtures and might be very toxic for the marine environments. One of the potential effects of the released water from the washing process is the depletion of dissolved oxygen levels in receiving waters (i.e., streams, canals, rivers, and sea) causing microbiological degradation. Although concerns on the potential ecological impact of these chemicals are growing, there is paucity of information on their ecotoxicity. Moreover, several studies have reported about microplastics in the marine environment, however, recent studies have reported the presence of microplastics also in the freshwater environments (Derraik 2002, Barboza 2019, Baldwin 2016). However, studies on the contribution of tetra-ethylene glycol and polyester to microbiological degradation and ecotoxicity are scarce.

Hence, our study aims to elucidate the chronic toxicity effects of polyester and tetra ethylene glycol (individually and in combination) in aquatic organisms. I employed a freshwater Daphnia magna, a very important organism that plays a very vital role in the freshwater food chain as a food source for many aquatic organisms. More importantly, the reason for using this organism as a test species is that it plays an important role in aquatic food chain, easy to breed, and it has a complete sequenced genome which is very advantageous for monitoring changes to their environment.

I produced microplastics from 100% polyester thread from Houjix, cut it finely and used a dissecting needle to cut into a size of 5mm or less through a microscope. I also included ethylene glycol to investigate its toxic effects on D. magna. Since my aim was also to compare the toxicity effect of both chemicals, I used different concentrations individually and then in combinations to determine the potential toxic effects of polyester and tetraethylene glycol (combined and separately) on the life (survival, death) of D. magna.

## 2. Materials and methods

### 2.1 Characteristic of D. magna

Daphnia magna plays an important role in water food chain and has sensitivity to temperature, resistance in a wide range of pH. Thus it is easy to breed and convenient to care for. Daphnia magna inhabits freshwater, adult length is between 1.5 and 5.0 mm, mean survivability is 56 days in 20 Celsius degrees. Daphnia magna becomes an adult in 7.5 days and starts first reproduction in 7~10 days after birth, repeating reproduction in 2~3 days.

### 2.2 Culture conditions for D. magna and algae

Culture medium used to grow the *D. magna* was prepared according to the following steps: took 20ml KCl (4g KCl/500ml of H2O), 20ml MgSO4 (60g MgSO4/500ml of H2O), 40ml NaHCO3 (48g Na-HCO3/500ml of H2O), and 100ml CaSO4.H2O (CaSO4.H2O/1000ml of H2O) from each solution and mixed them together. After mixing all the solutions, the distilled water was added to fill up the volume to 18.2 L. The solution was aerated for 24 hours.

To feed *D. magna*, the algae (Spirulina) were cultured according to the following steps: 1 ml of algae were inoculated in 100ml of sterile water and kept under the fluorescent light (25°C, 400ft/c). The intensity of light was 700 Lux to 800 Lux with 16 hr light and 8 hr dark photoperiod. The temperature to maintain the growth environment of algae was 18°C to 22°C.

### 2.3 Observation of population growth of D. magna

To observe the population growth of D.magna, I used the following criteria: observed population growth up to 3 generations through sub-culture, considered the adult size of *D. magna* as 1.5mm to 5mm, the polyester size was set to 0.5mm. Total of 100 D. magna individuals were group-cultured in a 2L beaker. 10 colonies were subcultured to another beaker. Subsequently, 100 young individuals were transferred into a fresh culture medium (2L beaker) and were colonized. This whole procedure was repeated continuously.

96 well plates were used in this experiment. Five neonates of *D. magna* were applied in a well and they were treated with the different concentrations of the solutions (TEG only, polyester only, TEG and polyester combination) to measure the IC50. Moreover, the microfiber size used in the study had a mean length of 2.96 × 10m (0.296mm). The temperature of the 96 well plate was maintained at 18°C to 22°C by immersing the cell plate in a container with fixed temperature. The individuals of *D. magna* were collected using a dropper with a diameter of 5mm or more and a disposable PVC dropper was used without rough edges. The heart rate of *D. magna* was measured with a counter watching the captured video with Samsung’s Galaxy S9+. Based on the heart rate, IC50 data is converted in the Prism software.

### 2.4 IC50 Measurement

360mg of 100% polyester yarn made by Japan’s Houjix, a thermostat, a water tank, a garden sieve, black sand, sand, a heater, mobile phone camera attached on microscope for heart rate measurement, a bubble generator, yeast powder, bottom filter, ring filter medium, thermometer, 330mg of perchloroethylene, optical microscope, mass, vial 30mlX10, beaker, distilled water, culture solution, and ultrapure water nitric acid were prepared to obtain the data. Polyester was cut to a certain size through a manual cutting method using an optical microscope and knife.

To check the lethal effects of different concentrations of TEG and polyester on *D. magna*, I used different concentrations (0mg/L, 4mg/L, 8mg/L, 16mg/L, 32mg/L, 64mg/L) of each solution. IC50 were measured to determine the half maximal inhibitory concentration and polyester on living *D. magna.* The exposure time ranged from 0 hour-24 hours (0 hr, 1 hr, 6 hrs, 12 hrs, 24 hrs) to count the heartbeat of *D. magna.*

Neonates who have not passed 24 hours since birth were used in the experiment. To minimize the negative impact on the results, I considered the following conditions. Due to the decreasing survival rate of the adult individuals, no more than 10 adults should be cultured in a 1L beaker to produce offspring that will be used in the results. Two hours prior to the test, the food was sufficiently provided. Limiting factors are observation time (exposure time of *D. magna* should be less than 1 minute under microscope), luminous Intensity and food (Not supplied before the measurement as mean heartbeat rate decreases lower than 10% after 1 hour of feeding). Collect *D. magna* with a dropper or pipette and transfer to a new culture medium. At this time, the diameter of the dropper or pipette should be at least 5mm larger than the adult Daphnia magna to minimize the physical stress of daphnia during movement.

The experiment was composed of four different groups. Group 1 was treated with different concentrations of TEG, group 2 was treated with different concentrations of polyester. In group 3, I used the combination of different concentrations of TEG and polyester. Group 4 consisted of D. magna without any treatment and was considered as a control.

## 3. Results and Discussion

The endpoints related to survival and growth in *D. magna* were not affected in the range of different concentrations of both tested compounds. After 24 hours of exposure to TEG and polyester, the number of D. magna was not reduced. I observed that the higher polyester concentrations adversely affected the population of *D. magna* with the death of neonates (20%). However, in the case of TEG treated *D. magna* neonates, there were no deaths. The control group exhibited no effects (data not shown).

This study showed that the IC50 of TEG was higher as compared to polyester which suggests that polyester was more adverse than TEG (Figure 1 and 2). Moreover, when TEG and polyester were treated in combination, IC50 value was lower (Figure 3) than the IC50 value of TEG and polyester. In other words, the TEG and polyester exhibited the lowest IC50 value in this study. These results suggest that TEG and polyester in combination had adverse effects on the growth of *D. magna* (Jang 2015).

**figure 1.**
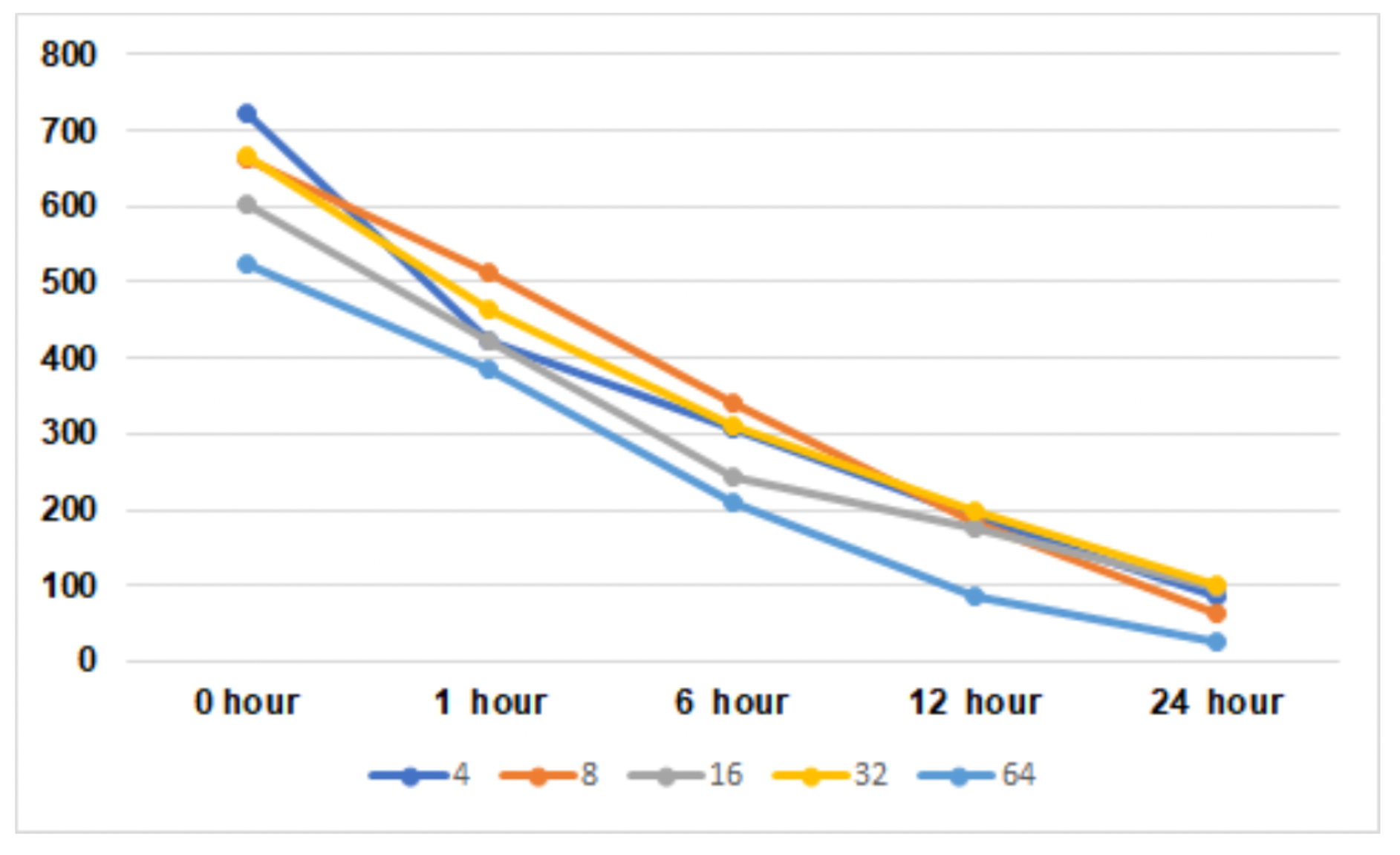
only polyester treated on *D. magna*

**figure 2.**
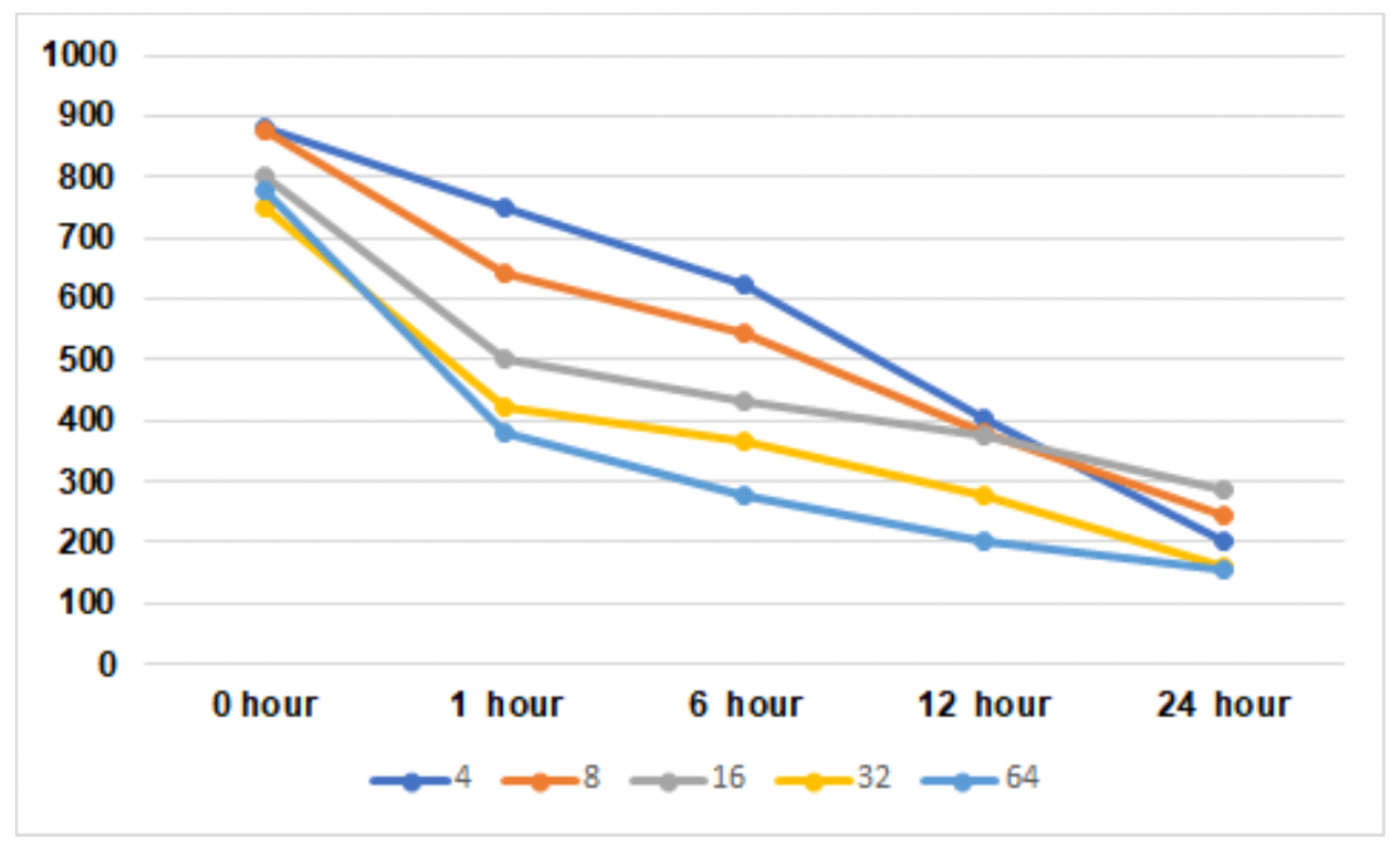
only TEG treated on *D. magna*

**figure 3.**
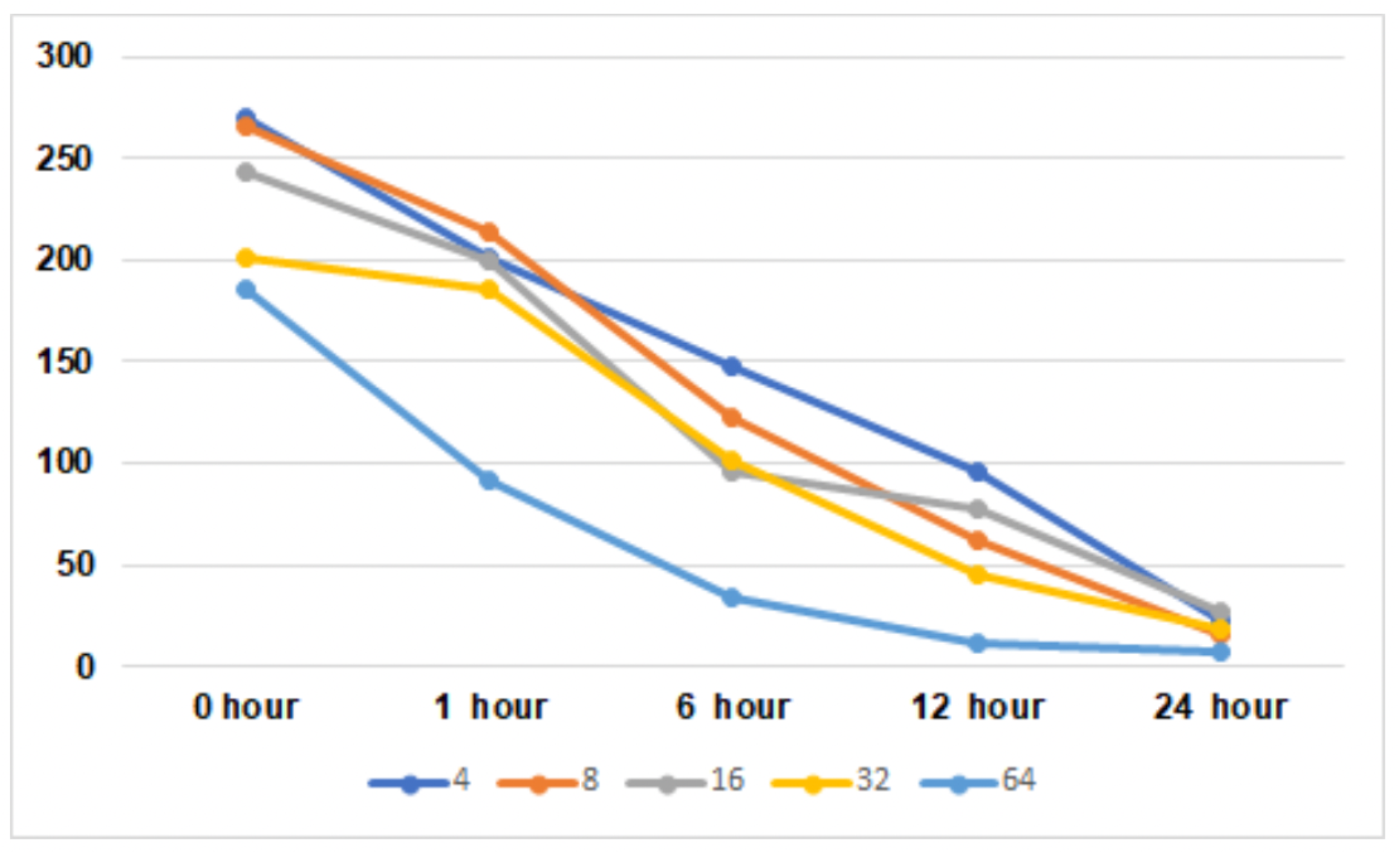
polyester and TEG treated on *D. magna*

This study suggest that a combined solution of TEG and polyester could have adverse effects on the growth and development of D. magna and even may cause the death.

The higher the concentration, the lower the IC50 value at the same time period, and it gradually decreased as time passed.

As the same trend that had shown in figure 1, IC50 decreased as time passed, and only the TEG treated group showed a larger range of IC50 compared to only polyester treated group. Thus, I can say that the impact of polyester on D. magna is greater than TEG.

IC50 of polyester and TEG treated group shows a much larger range of IC50 values than the TEG only treated, and polyester only treated groups. At 32mg/L, the most irregular IC50 value changes, and at 4mg/L, 8mg/L, and 16mg/L, it shows a relatively regular decrease. Note that there is a significant decrease in IC50 values at 32 mg/L.

In this study I investigated that the combined solution of polyester and TEG impacts on *D. magna* greater than other solutions. Significantly, in case of the combined solution, *D. magna* mostly died when the experiment ended at the same time few D. magna survived after the experiment in other solutions. This indicates the toxicity of the combined solution is high enough to affect harmness on *D. magna.* Based on this study, the effect of microsynthetic fibers and organic solvents on the environment on the population of Daphnia, the primary consumer of freshwater aquatic ecosystems, can be investigated. Finding ways to replace or eliminate consumption of fine synthetic fibers and organic solvents in real life as a way to solve these problems by identifying the factors of a specific ecosystem that are affected by fine synthetic fibers and organic solvents, which account for a significant portion of environmental pollutants If so, it can help to restore the natural ecosystem.

